# ExpOmics: a comprehensive web platform empowering biologists with robust multi-omics data analysis capabilities

**DOI:** 10.1101/2024.04.23.588859

**Authors:** Douyue Li, Zhuochao Min, Jia Guo, Yubin Chen, Wenliang Zhang

## Abstract

**Motivation:** High-throughput technologies have yielded a broad spectrum of multi-omics datasets, offering unparalleled insights into complex biological systems. However, effectively analyzing this diverse array of data presents challenges, given factors such as species diversity, data types, costs, and limitations of available tools.

**Results:** We propose ExpOmics, a comprehensive web platform featuring seven applications and four toolkits with 28 customizable analysis functions, spanning various aspects of differential expression, co-expression, WGCNA analysis, feature selection, and functional enrichment analysis. ExpOmics empowers users to effortlessly upload and explore multi-omics data without organism restriction, supporting a wide array of data types including gene, mRNA, lncRNA, miRNA, circRNA, piRNA, and protein expression data. It is compatible with diverse gene nomenclatures and expression value types. Moreover, ExpOmics enables users to comprehensive analysis of 22,427 transcriptomic datasets sourced from 63 projects and 196 cancer subtypes in TCGA, discovering cancer biomarkers and targets. The analysis results from ExpOmics are visually presented in high-quality graphical formats suitable for publication, available for free download. In summary, ExpOmics can serves as a robust platform for global researchers to delve into diverse expression datasets, gain biological insights, and formulate testable hypotheses.

**Availability and implementation:** ExpOmics is available at http://www.biomedical-web.com/expomics.

## 1 Introduction

Multi-omics data, generated by high-throughput technologies like next-generation sequencing and mass spectrometry, are invaluable resources for understanding complex biological systems (Stark et al. 2019; Wang et al. 2009). However, despite their potential, mining and analyzing such data remain challenging due to high costs and thresholds. For example, the complexity and diversity of of multi-omics datasets presents a significant challenge for biologists lacking bioinformatics expertise to explore these data, including gene, mRNA, lncRNA, miRNA, circRNA, piRNA, and protein expression data (Mougin et al. 2018). Furthermore, customizable data analysis is often required to address specific biological questions, emphasizing the necessity for user-friendly web platforms to enable efficient investigation of these complex and diverse data and advance life science research (Mougin, et al. 2018; O’Donoghue et al. 2010). To tackle these challenges, bioinformaticians have made efforts and developed several web platforms to explore and analyze multi-omics data (Chang et al. 2018; Cheng et al. 2021; Conard et al. 2021; Ge et al. 2018; Liu et al. 2022; Zhou et al. 2021).

However, there still remain some critical issues that urgently need to be addressed. For example, ExpressVis (Liu, et al. 2022), eVITTA (Cheng, et al. 2021), PANDA-view (Chang, et al. 2018), iDEP (Ge, et al. 2018), and TIMEOR (Conard, et al. 2021) primarily focus on gene and protein expression data, catering to specific species like Homo sapiens and model organisms. Consequently, they lack the capability to handle non-coding gene expression data and data from other organisms, including lncRNA, circRNA, miRNA, and piRNA. Moreover, they lack essential analysis capabilities like weighted gene co-expression network analysis (WGCNA) and feature selection, as well as integration with large-scale international omics projects such as The Cancer Genome Atlas (TCGA) (Weinstein et al. 2013). Furthermore, they provide relatively limited functionality for differential expression and functional enrichment analysis, which cannot meet the demands of complex analyses.

To fill these gaps, we propose ExpOmics, an easy-to-use web platform featuring applications designed to assist biologists in efficiently processing various types of expression data without requiring programming skills. ExpOmics offers robust multi-omics data analysis and visualization capabilities for exploring gene, mRNA/lncRNA, miRNA, circRNA, piRNA, and protein expression data, covering various aspects of differential expression, co-expression, WGCNA analysis, feature selection, and functional enrichment analysis. ExpOmics supports users to upload the expression data with diverse gene nomenclatures from different resources and support various types of expression values and any organisms. Additionally, ExpOmics have integrated 22,427 transcriptomic datasets of 196 cancer subtypes from TCGA. This integration empowers biologists to thoroughly explore these data, facilitating the discovery and validation of cancer biomarkers and targets. In summary, ExpOmics serves as a valuable web platform for empowering biologists worldwide with robust multi-omics data analysis capabilities.

## 2 Materials and methods

### 2.1 Implementation of ExpOmics

The ExpOmics web platform was developed using a front-end and back-end separation framework, which was previously described (Zhang et al. 2022; Zhang et al. 2022). Within ExpOmics, seven distinct applications (including GeneExplyzer, Transcriptlyzer, miRExplyzer, circExplyzer, piRExplyzer, ProteinExplyzer, and TCGAExplyzer) with four toolkits (including DiffExpToolkit, CorrExpToolkit, WGCNAToolkit, and FeatureSelectToolkit) were developed based on the R project (Version 4.3.2) (Figure 1 and Table 1), seamlessly integrated to provide a comprehensive suite of functionalities (Table 2). The implementation of these applications and toolkits relies on a carefully selected set of R packages and resources, meticulously detailed in Table 1, Table S1, and Table S2.

**Figure 1.**
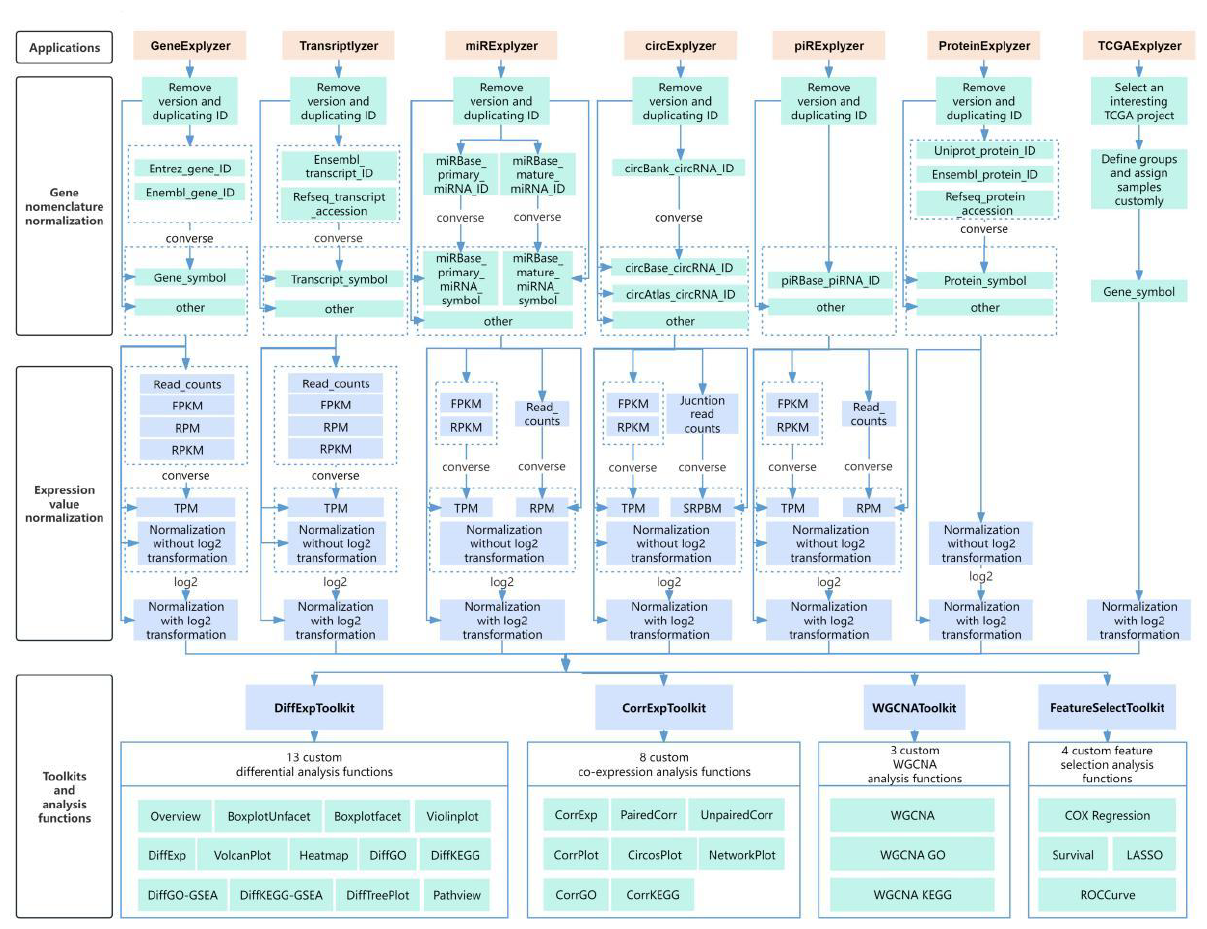
The implementation of ExpOmics.

**Table 1.**
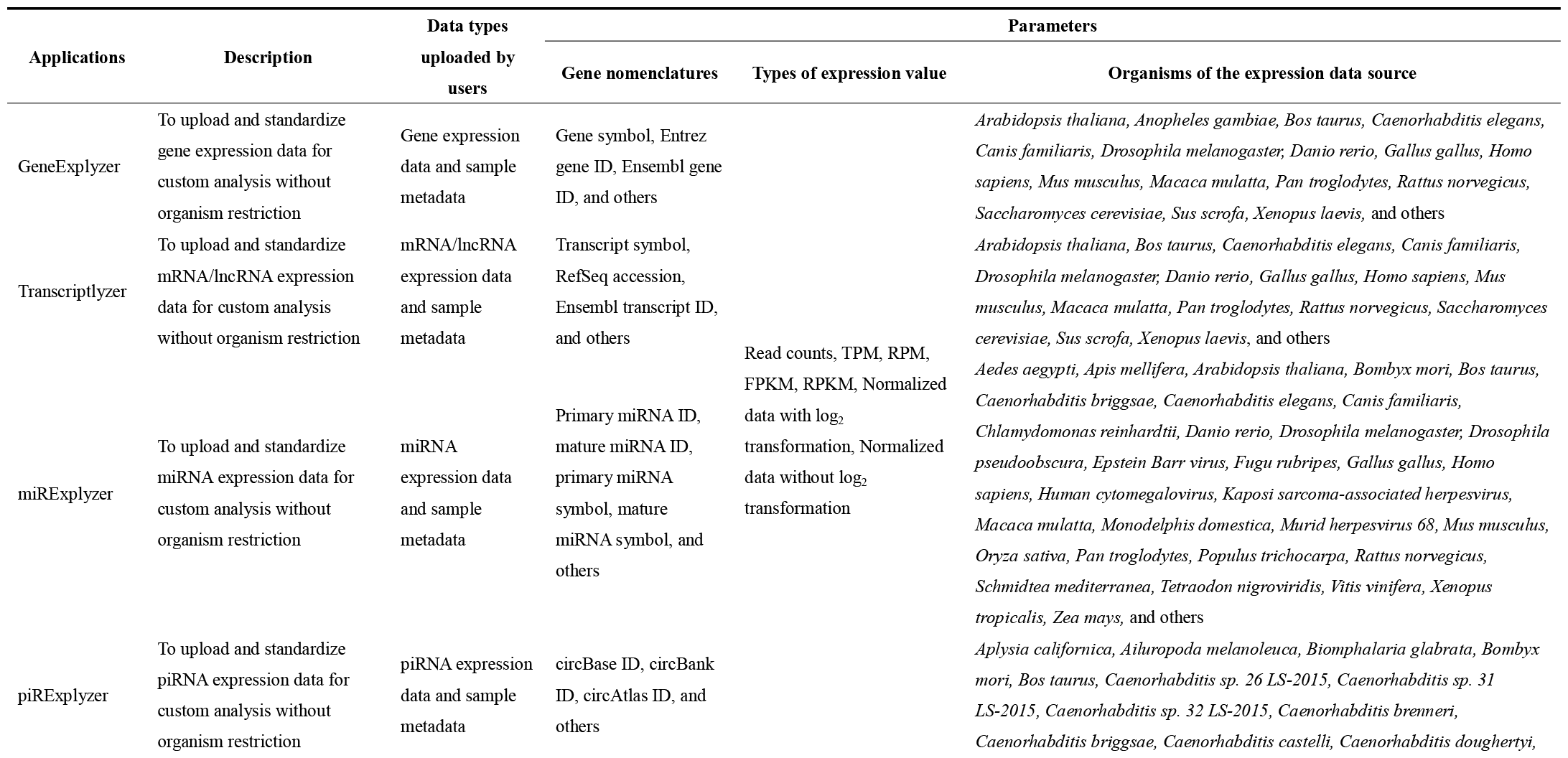
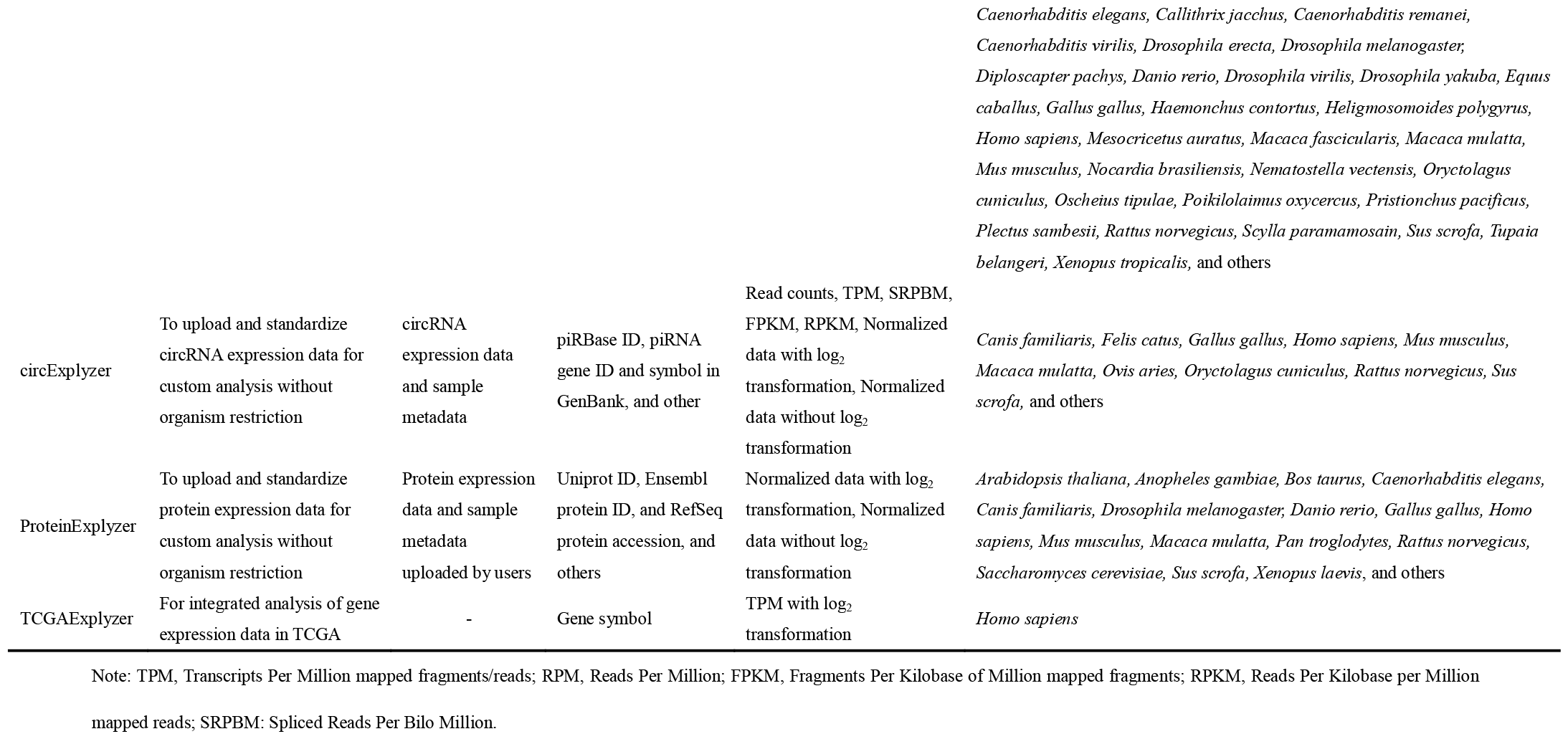
The details of the implementation and utilization of the seven applications in ExpOmics.

**Table 2.**
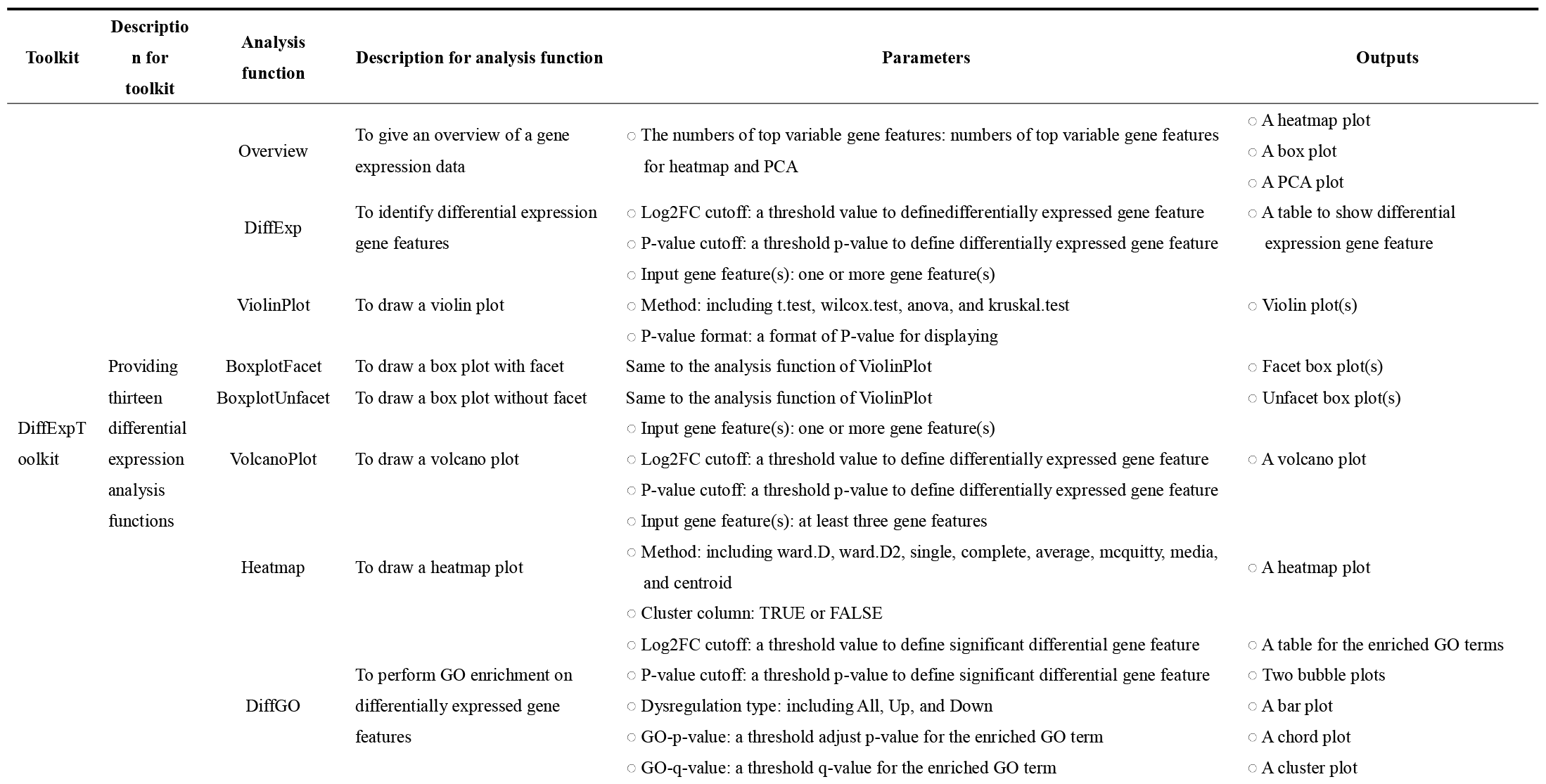
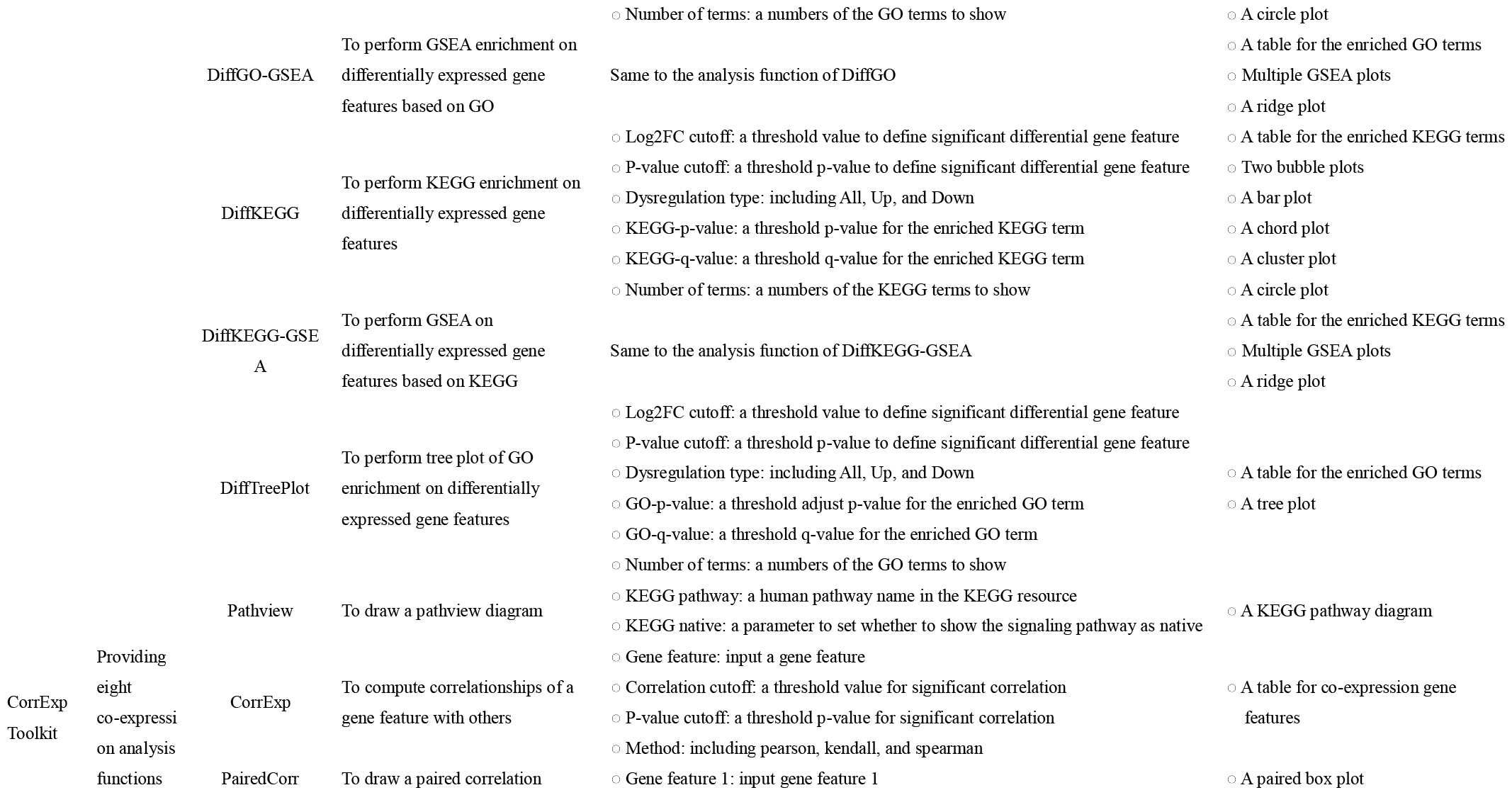
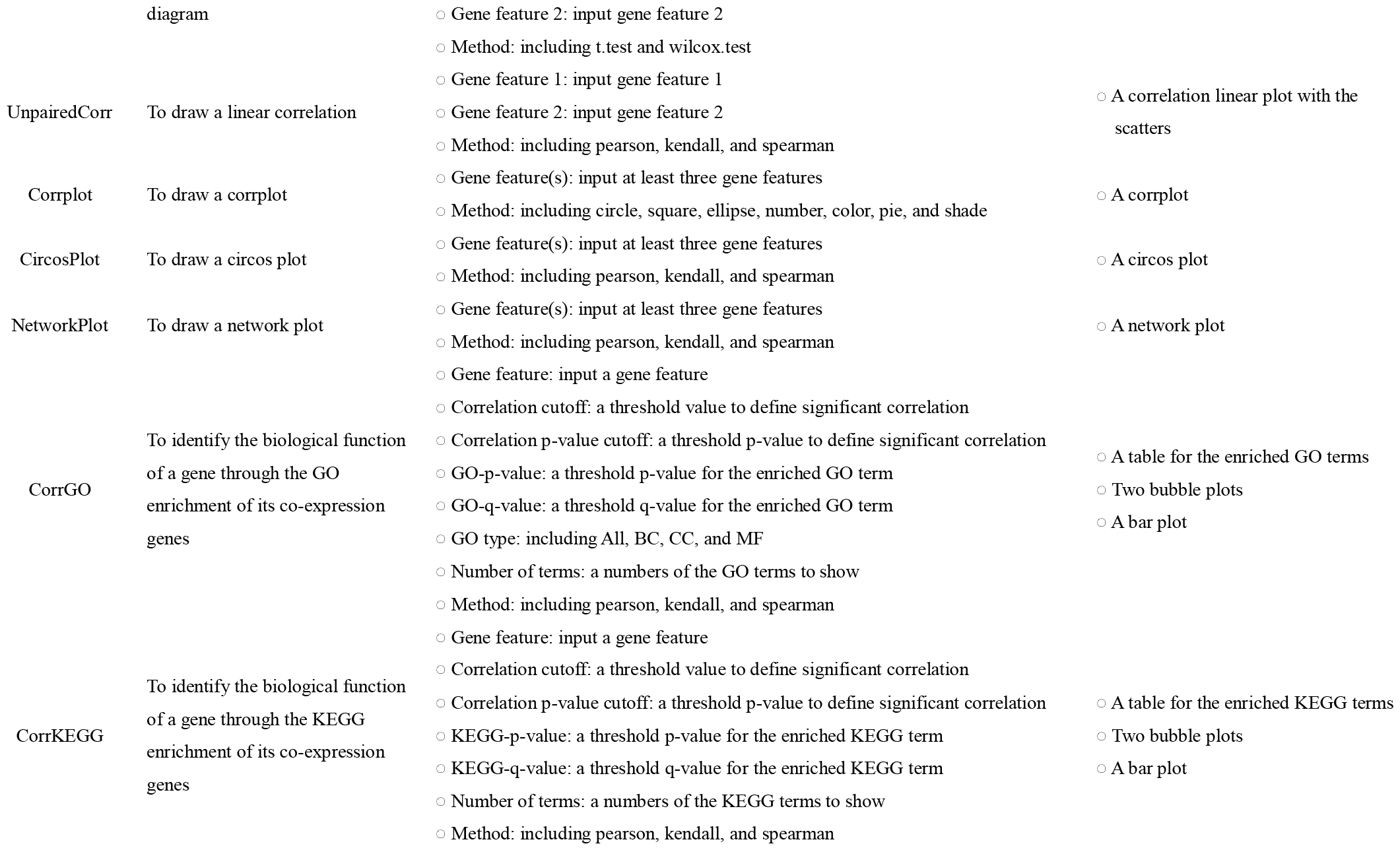
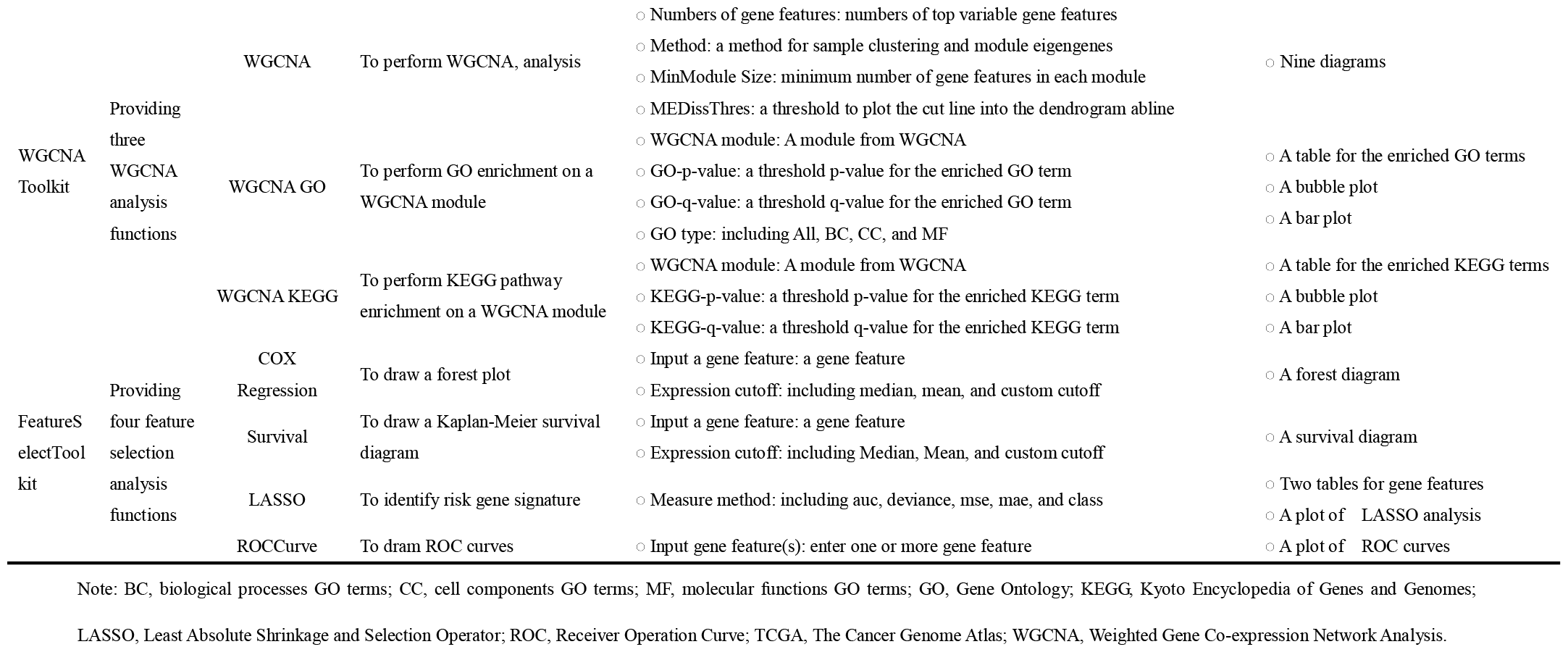
The details of the implementation and utilization of the four toolkit in ExpOmics.

### 2.3 Implementation of miRExplyzer

To implement miRExplyzer, miRNA annotations spanning 31 organisms were extracted from the miRBase database (Kozomara et al. 2019). These annotations include primary miRNA ID, mature miRNA ID, primary miRNA symbol, mature miRNA symbol, and other relevant details. Subsequently, utilizing the extracted miRNA annotations, we developed miRExplyzer based on the R project (Figure 1, Table 1, and Table S1).

### 2.4 Implementation of circExplyzer

For circExplyzer, circRNA annotations spanning 10 organisms were extracted from databases such as circBase (Glazar et al. 2014), circBank (Liu et al. 2019), and circAtlas (Wu et al. 2024). These annotations include circRNA ID, host gene annotations, and other pertinent details. Leveraging this information, circExplyzer was meticulously developed based on the R project (Figure 1, Table 1, and Table S1).

### 2.5 Implementation of piRExplyzer

To implement piRExplyzer, piRNA annotations spanning 43 organisms were extracted from piRBase (Wang et al. 2022). These annotations include piRNA ID, piRNA gene ID and symbol in GenBank, along with other relevant details. Utilizing the extracted piRNA annotations, we developed piRExplyzer based on the R project (Figure 1, Table 1, and Table S1).

### 2.6 Implementation of ProteinExplyzer

For ProteinExplyzer, protein annotations of 16 organisms were extracted from Uniprot (2023). These annotations include protein ID and symbol in databases such as Uniprot, Ensembl, and RefSeq, along with their host gene and other relevant details. Leveraging this information, ProteinExplyzer was developed based on the R project (Figure 1, Table 1, and Table S1).

### 2.7 Implementation of TCGAExplyzer

To implement TCGAExplyzer, we have extracted and curated gene expression data and sample metadata sourced from TCGA, encompassing 63 projects and 22,427 transcriptomics datasets across 196 cancer subtypes via the GDC portal (Weinstein, et al. 2013) (Table 1). Additionally, four toolkits featuring 28 analysis functions were developed and incorporated into TCGAExplyzer based on the R project (Figure 1, Table 1, and Table S1).

### 2.8 Implementation of DiffExpToolkit, CorrExpToolkit, WGCNAToolkit, and FeatureSelectToolkit

The toolkits, were designed to provide 28 analysis functions for various aspects of differential expression, co-expression, WGCNA, and feature selection analysis, including DiffExpToolkit, CorrExpToolkit, WGCNAToolkit, and FeatureSelectToolkit (Figure 1 and Table 2). These analysis functions are designed for enabling users to select a project, assign samples to predefined groups, and set parameters for custom analysis. The R packages and resources used to implement these toolkits and analysis functions are detailed in Table S2.

## 3 Results

### 3.1 Overview of ExpOmics

ExpOmics (http://www.biomedical-web.com/expomics) is a user-friendly web platform accessible for free without requiring login. It offers seven meticulously designed applications: GeneExplyzer, Transcriptlyzer, miRExplyzer, circExplyzer, piRExplyze, ProteinExplyzer, and TCGAExplyzer (Figure 2A and Table 1). Within these applications, the applications of GeneExplyzer, Transcriptlyzer, miRExplyzer, circExplyzer, piRExplyze, and ProteinExplyzer allow users to upload, standardize, and explore expression data of gene, mRNA, lncRNA, miRNA, circRNA, piRNA, and protein originating from various organisms (Table 1). Users can specify three parameters: organism, gene nomenclature, and expression value type, enabling standardization and analysis across different species with varying gene nomenclatures and expression value types (Figure 2A).

**Figure 2.**
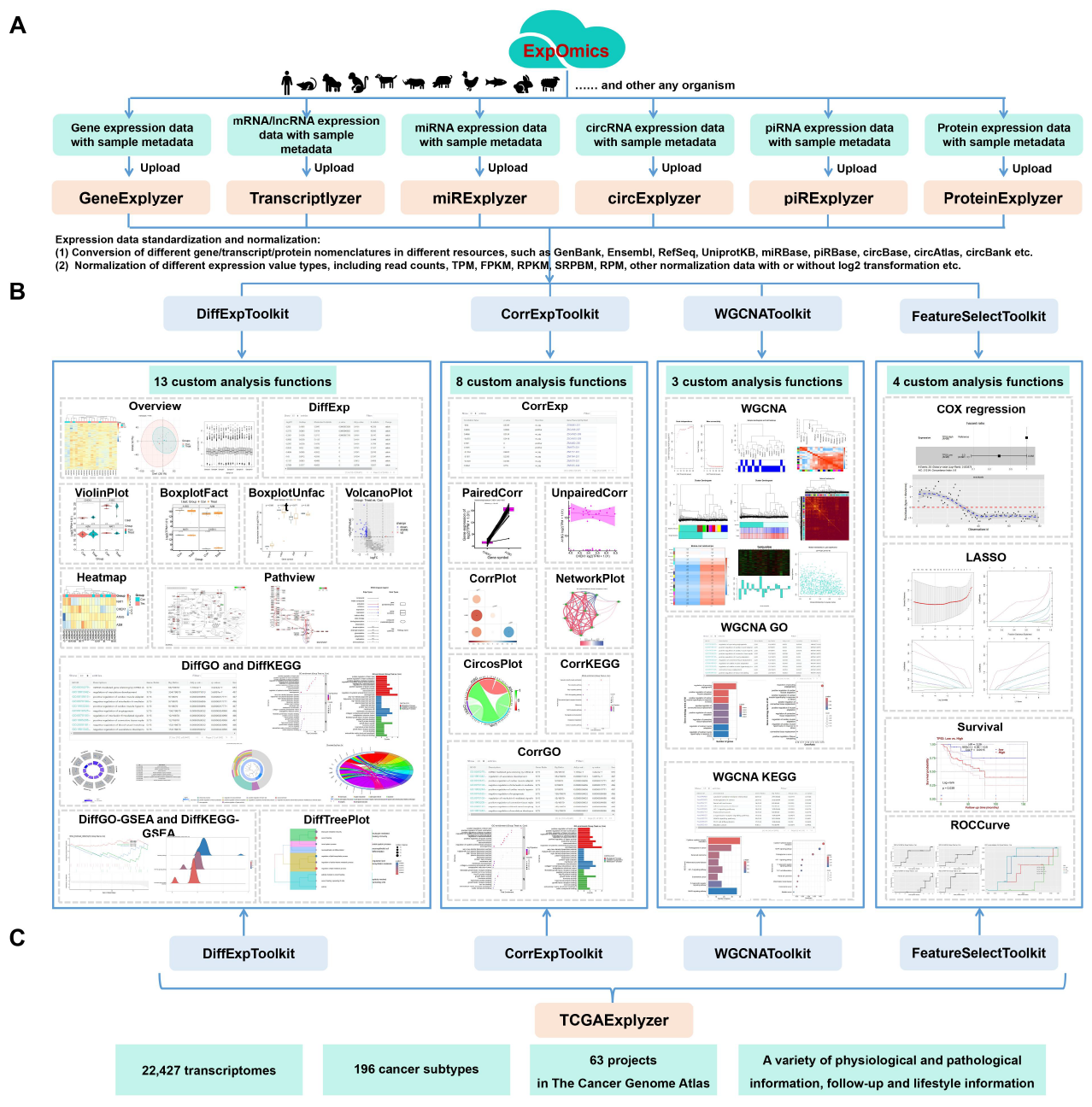
The applications, toolkits, and analysis functions in ExpOmics.

Additionally, users can customize organism and gene nomenclature parameters to “other” for scenarios not covered by default options. Upon completion of data uploading and standardization, the standardized expression data is stored and assigned a unique identifier for tracking and subsequent analysis. ExpOmics provides a feature called data remove for users to delete their uploaded data using the assigned unique identifier, ensuring data security. Each application were equipped with four toolkits with 28 analysis functions for comprehensive exploration of gene, mRNA, lncRNA, miRNA, circRNA, piRNA, and protein expression data across various organisms (Figure 2B, Table 1 and 2). These toolkits include DiffExpToolkit, CorrExpToolkit, WGCNAToolkit, and FeatureSelectToolkit. Moreover, all of these analysis functions enable users to select a project, assign samples to predefined groups, and set parameters for custom analysis based on the sample metadata of the dataset.

In contrast to the GeneExplyzer, Transcriptlyzer, miRExplyzer, circExplyzer, piRExplyze, and ProteinExplyzer applications, TCGAExplyzer focuses on comprehensive and custom analysis of 22,427 transcriptomic datasets of 63 projects and 196 cancer subtypes in TCGA (Figure 2C). Furthermore, the results of these analysis functions are visually presented in high-quality graphical formats suitable for publication, available for free download. Furthermore, the result tables are equipped with filtering and sorting capabilities, facilitating users in effortlessly exploring data of interest.

Detailed user guidance and video tutorials are available on the “Help” page and individual application webpages. Extensive tests were performed across popular web browsers (Google Chrome, Safari, Microsoft Edge, and Firefox) to ensure optimal performance. Details of these applications and toolkits are detailed below.

### 3.2 Seven applications

The seven applications, namely GeneExplyzer, miRExplyzer, circExplyzer, piRExplyzer, ProteinExplyzer, and TCGAExplyzer, have been described below. Their detailed illustrations can be found in Figure 1, Figure 2, and Table 1.

#### 3.2.1 GeneExplyzer

GeneExplyzer aims to remove duplicate genes or mRNA/lncRNAs and convert them into standardized gene or transcript symbols in user-uploaded expression data, applicable to listed 13 organisms and others. Moreover, it adeptly normalizes different expression values, including read counts, Reads Per Kilobase per Million mapped reads (RPKM), and Fragments Per Kilobase of exon model per Million mapped fragments (FPKM), into Transcripts Per Million (TPM), while seamlessly integrating a log2 transformation process for enhanced analysis.

#### 3.2.2 miRExplyzer

miRExplyzer aims to remove duplicate miRNA IDs and convert theminto their standardized miRNA symbols in user-uploaded expression data, applicable to listed 31 organisms and others. Moreover, miRExplyzer adeptly normalizes different expression values, such as normalizing read counts to Reads Per Million (RPM) with a log2 transformation and adjusting RPKM and FPKM to TPM with a log2 transformation.

#### 3.2.3 circExplyzer

circExplyzer aims to remove duplicate circRNAs and convert them into their standardized circRNA IDs in user-uploaded expression data, applicable to listed 10 organisms and others. Additionally, circExplyzer adeptly normalizes different expression values, such as normalizing junction read counts to SRPBM with a log2 transformation and adjusting RPKM and FPKM to TPM with a log2 transformation.

#### 3.2.4 piRExplyzer

piRExplyzer aims to remove duplicate piRNAs and convert them into their standardized piRNA IDs in user-uploaded expression data, applicable to listed 43 organisms and others . Additionally, piRExplyzer adeptly normalizes diverse expression values, such as read counts to RPM with a log2 transformation and RPKM and FPKM to TPM with a log2 transformation.

#### 3.2.5 ProteinExplyzer

ProteinExplyzer aims to remove duplicate proteins and convert them into their standardized protein IDs combined with their host gene symbol in user-uploaded expression data for easier recognition, applicable to listed 16 organisms and others. Additionally, ProteinExplyzer adeptly applies a log2 transformation to the uploaded expression data if the transformation was not initially applied.

#### 3.2.6 TCGAExplyzer

TCGAExplyzer hosts 22,427 transcriptomic datasets of 63 projects and 196 cancer subtypes in TCGA. Moreover, TCGAExplyzer provides four toolkits with the 28 custom analysis functions for comprehensive analysis of these data to expedite the discovery of cancer biomarkers and targets.

### 3.3 Four toolkits

The four applications, namely DiffExpToolkit, CorrExpToolkit, WGCNAToolkit, and FeatureSelectToolkit, have been described below. Their detailed illustrations can be found in Figure 1, Figure 2, and Table 2.

#### 3.3.1 DiffExpToolkit

The DiffExpToolkit offers 13 analysis functions for conducting various aspects of differential expression analysis and visualization on the gene, mRNA, lncRNA, miRNA, circRNA, piRNA, and protein expression data from different organisms. These functions include Overview, DiffExp, ViolinPlot, BoxplotFacet, BoxplotUnfacet, VolcanoPlot, Heatmap, DiffGO, DiffGO-GSEA, DiffKEGG, DiffKEGG-GSEA, DiffTreePlot, and Pathview.

#### 3.3.2 CorrExpToolkit

The CorrExpToolkit provides 8 co-expression analysis functions for conducting various aspects of co-expression analysis and visualization on the gene, mRNA, lncRNA, miRNA, circRNA, piRNA, and protein expression data from different organisms. These functions encompass CorrExp, PairedCorr, UnpairedCorr, CorrPlot, CircosPlot, NetworkPlot, CorrGO, and CorrKEGG.

#### 3.3.3 WGCNAToolkit

WGCNAToolkit enables users to conduct WGCNA analysis, including network construction, module detection, gene selection, topological property calculations, data simulation, and GO and KEGG pathway enrichment analysis on the gene, mRNA, lncRNA, miRNA, circRNA, piRNA, and protein expression data from different organisms.

#### 3.3.4 FeatureSelectToolkit

FeatureSelectToolkit provides 4 analysis functions for conducting various aspects of feature selection analysis on the gene, mRNA, lncRNA, miRNA, circRNA, piRNA, and protein expression data from different organisms, identifying and discovering important gene features. These functions include COX regression, LASSO, ROCCurve, and Survival.

## 4 Discussion

ExpOmics is a easy-to-use web platform that provides powerful multi-omics data analysis capabilities for biologists. Compared to existing web platforms, ExpOmics has a wider range of applications and stronger analysis capabilities (Table 3). For example, compared to mainstream multi-omics web platforms like ExpressVis (Liu, et al. 2022), eVITTA (Cheng, et al. 2021), and OmicsAnalyst (Zhou, et al. 2021), ExpOmics not only supports the expression data of genes, mRNA, and proteins as they do, but also additionally supports the analysis of non-coding RNA expression data, including lncRNA, circRNA, miRNA, and piRNA etc. Moreover, ExpOmics is capable of accommodating expression data without species restrictions, whereas ExpressVis and eVITTA only support data from model organisms like *Homo sapiens, Mus musculus, Rattus norvegicus etc*. Additionally, although OmicsAnalyst (Zhou, et al. 2021) can handle expression data without species restrictions, it lacks the capacity to perform functional enrichment. ExpressVis, eVITTA, and OmicsAnalyst also lack WGCNA analysis functionality, whereas ExpOmics specifically provides this feature. WGCNA analysis can unveil intricate relationships among diverse gene features and elucidate how these relationships correlate with phenotypic traits (Langfelder and Horvath 2008). Compared to ExpOmics, their capabilities for both differential expression analysis and co-expression analysis are limited, and they have not developed functional modules for the integrated analysis of TCGA data. Taken together, ExpOmics demonstrates superior performance and more comprehensive functionality compared to existing similar web platforms.

**Table 3.**
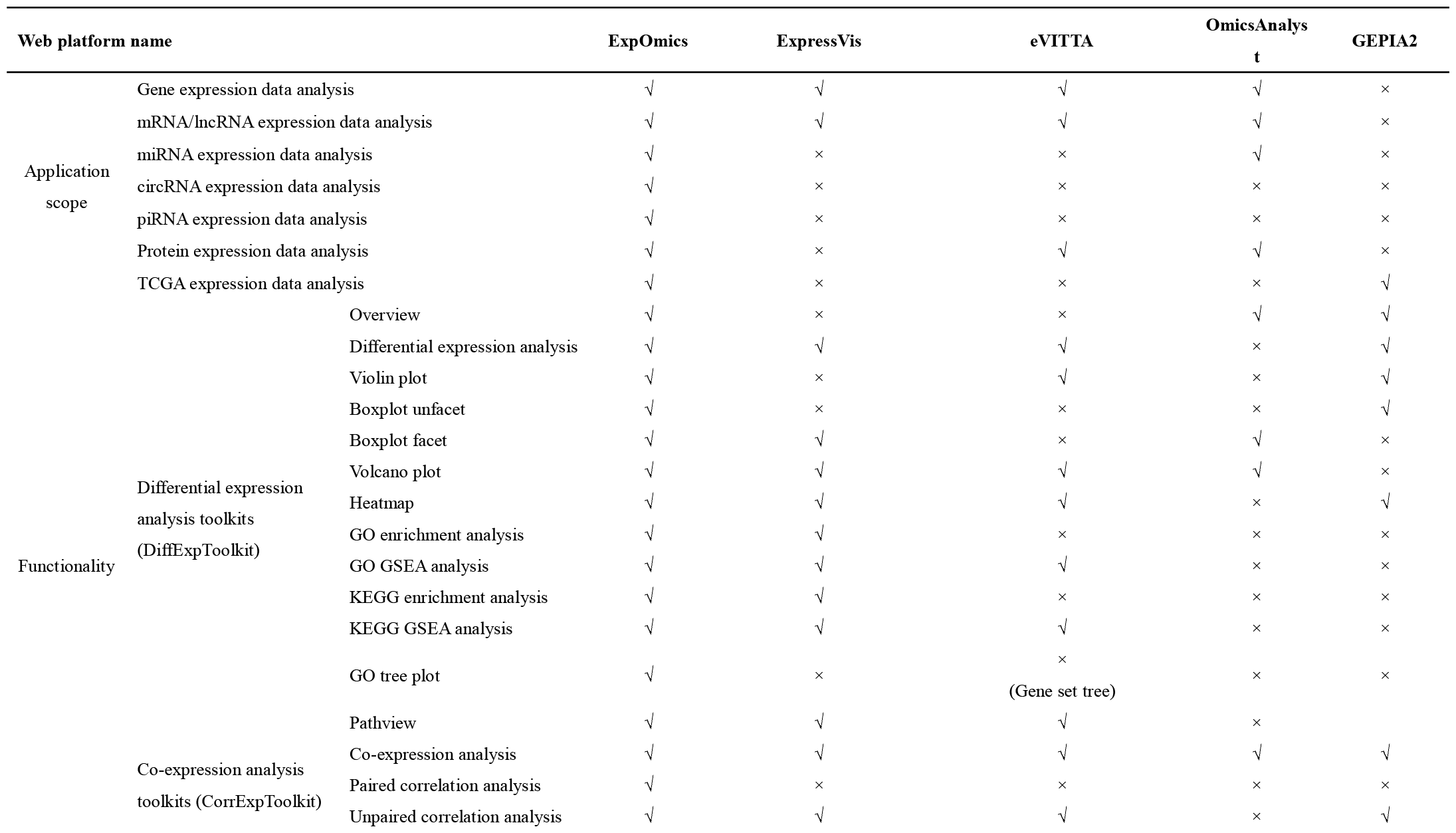
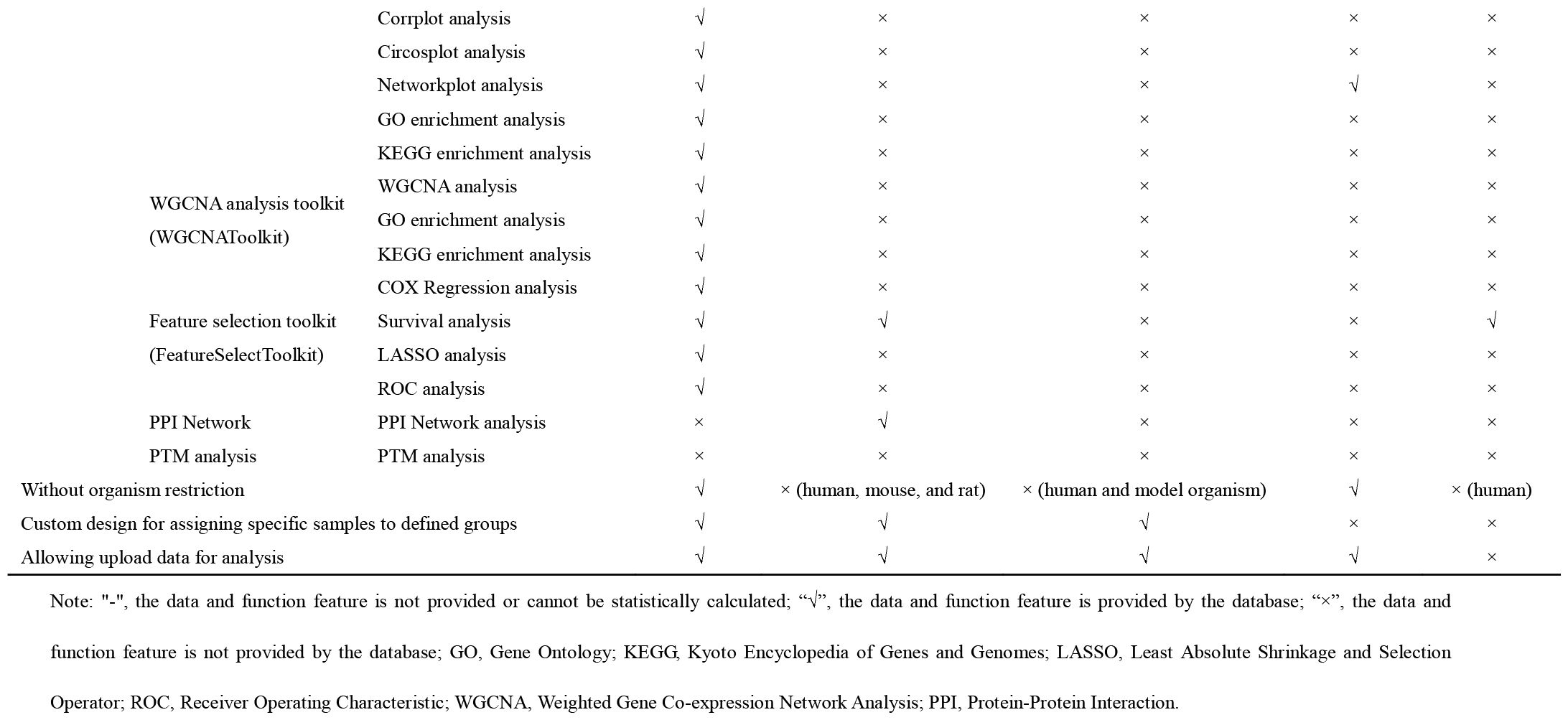
The comparison of application scope and functionality of ExpOmics with other web platform.

## 5 Future plans

In the foreseeable future, we plan to enhance ProteinExplyzer in ExpOmics by integrating a Protein-Protein Interaction (PPI) network analysis function. And the web platform will be updated to be capable of adapting expression data at larger scales, like single cell and spatial transcriptome, and more custom parameters, like plot size, plot color scheme, user-defined Gene Matrix Transposed (gmt) file and Gene Expression Omnibus Platform file. We are committed to the ongoing maintenance and improvement of ExpOmics at analytical and visualization capabilities, and expect these updates will significantly boost the utility of ExpOmics in the analysis of multi-omics data.

## Data availability

ExpOmics is available at http://www.biomedical-web.com/expomics. The web platform is freely accessible without login requirement.

## References

Chang C, Xu K, Guo C et al. PANDA-view: an easy-to-use tool for statistical analysis and visualization of quantitative proteomics data. Bioinformatics 2018;34: 3594–6.

Cheng X, Yan J, Liu Y et al. eVITTA: a web-based visualization and inference toolbox for transcriptome analysis. Nucleic Acids Res 2021;49: W207–15.

Conard A M, Goodman N, Hu Y et al. TIMEOR: a web-based tool to uncover temporal regulatory mechanisms from multi-omics data. Nucleic Acids Res 2021;49: W641–53.

Ge S X, Son E W, Yao R iDEP: an integrated web application for differential expression and pathway analysis of RNA-Seq data. Bmc Bioinformatics 2018;19: 534.

Glazar P, Papavasileiou P, Rajewsky N circBase: a database for circular RNAs. Rna 2014;20: 1666–70.

Kozomara A, Birgaoanu M, Griffiths-Jones S miRBase: from microRNA sequences to function. Nucleic Acids Res 2019;47: D155–62.

Langfelder P, Horvath S WGCNA: an R package for weighted correlation network analysis. Bmc Bioinformatics 2008;9: 559.

Liu M, Wang Q, Shen J et al. Circbank: a comprehensive database for circRNA with standard nomenclature. Rna Biol 2019;16: 899–905.

Liu X, Xu K, Tao X et al. ExpressVis: a biologist-oriented interactive web platform for exploring multi-omics data. Nucleic Acids Res 2022;50: W312–21.

Mougin F, Auber D, Bourqui R et al. Visualizing omics and clinical data: Which challenges for dealing with their variety? Methods 2018;132: 3–18.

O’Donoghue S I, Gavin A C, Gehlenborg N et al. Visualizing biological data-now and in the future. Nat Methods 2010;7: S2–4.

Sayers E W, Cavanaugh M, Clark K et al. GenBank 2024 Update. Nucleic Acids Res 2024;52: D134–7.

Stark R, Grzelak M, Hadfield J RNA sequencing: the teenage years. Nat Rev Genet 2019;20: 631–56.

UniProt Consortium. UniProt: the Universal Protein Knowledgebase in 2023. Nucleic Acids Res 2023;51: D523–31.

Wang J, Shi Y, Zhou H et al. piRBase: integrating piRNA annotation in all aspects. Nucleic Acids Res 2022;50: D265–72.

Wang Z, Gerstein M, Snyder M RNA-Seq: a revolutionary tool for transcriptomics. Nat Rev Genet 2009;10: 57–63.

Weinstein J N, Collisson E A, Mills G B et al./person-group>. The Cancer Genome Atlas Pan-Cancer analysis project. Nat Genet 2013;45: 1113–20.

Wu W, Zhao F, Zhang J circAtlas 3.0: a gateway to 3 million curated vertebrate circular RNAs based on a standardized nomenclature scheme. Nucleic Acids Res 2024;52: D52–60.

Zhang W, Liu Y, Min Z et al. circMine: a comprehensive database to integrate, analyze and visualize human disease-related circRNA transcriptome. Nucleic Acids Res 2022;50: D83–92.

Zhang W, Zhang Y, Min Z et al. COVID19db: a comprehensive database platform to discover potential drugs and targets of COVID-19 at whole transcriptomic scale. Nucleic Acids Res 2022;50: D747–57.

Zhou G, Ewald J, Xia J OmicsAnalyst: a comprehensive web-based platform for visual analytics of multi-omics data. Nucleic Acids Res 2021;49: W476–82.

