## Supplementary material for "ExpOmics: a comprehensive web platform empowering biologists with robust multi-omics data analysis capabilities": Table S1

**Table S1. The details of the implementation of the four toolkit in ExpOmics**

| **Application** | **Resources and R packages** |
| --- | --- |
| GeneExplyzer | ENSEMBL v2023, GENCODE v2023, NCBI GenBank v2023, clusterProfiler v3.18.1, data.table v1.14.0, dplyr v1.0.7, jsonlite v1.7.2, stringr v1.4.0, orgDbs (like org.Hs.eg.db v3.16.0) for the listed organisms, tidyverse v1.3.1 |
| Transcriptlyzer | ENSEMBL v2023, GENCODE v2023, NCBI GenBank v2023, data.table v1.14.0, dplyr v1.0.7, jsonlite v1.7.2, stringr v1.4.0, tidyverse v1.3.1 |
| miRExplyzer | miRBase v22.1, miEAA v2.1, data.table v1.14.0, dplyr v1.0.7, jsonlite v1.7.2, stringr v1.4.0, tidyverse v1.3.1 |
| piRExplyzer | piRBase v3.0, data.table v1.14.0, dplyr v1.0.7, jsonlite v1.7.2, stringr v1.4.0, tidyverse v1.3.1 |
| circExplyzer | circAtlas v3.0, circBank v2019, circBase v2014, data.table v1.14.0, dplyr v1.0.7, jsonlite v1.7.2, stringr v1.4.0, tidyverse v1.3.1 |
| ProteinExplyzer | UNIPROT v2023, ENSEMBL v2023, NCBI gene info v2023,  data.table v1.14.0, dplyr v1.0.7, jsonlite v1.7.2, stringr v1.4.0, tidyverse v1.3.1 |
| TCGAExplyzer | The Cancer Genome Atlas (TCGA) v2023 |
