## Supplementary material for "ExpOmics: a comprehensive web platform empowering biologists with robust multi-omics data analysis capabilities": Table S2

**Table S2. The details of the implementation of the four toolkit in ExpOmics**

| **Toolkit** | **Analysis function** | **Resources and R packages** |
| --- | --- | --- |
| DiffExpToolkit | Overview | dplyr v1.0.7, factoextra v1.0.7, FactoMine v1.08, ggplot2 v3.3.5, pheatmap v1.0.12 |
|  | DiffExp | dplyr v1.0.7, jsonlite v1.7.2, limma v3.58.1 |
|  | ViolinPlot | dplyr v1.0.7, easyGgplot2 v1.0.0.9000, EnvStats v2.8.1, ggplot2 v3.3.5, ggpubr v0.4.0, reshape2 v1.4.4, usethis v2.2.2 |
|  | BoxplotFacet | dplyr v1.0.7, ggplot2 v3.3.5, ggpubr v0.4.0, gridExtra v2.3, reshape2 v1.4.4 |
|  | BoxplotUnfacet | Same to the analysis function of BoxplotFacet |
|  | VolcanoPlot | dplyr v1.0.7, ggplot2 v3.3.5, limma v3.58.1 |
|  | Heatmap | dplyr v1.0.7, ggplot2 v3.3.5, jsonlite v1.7.2, pheatmap v1.0.12 |
|  | DiffGO | GO database, clusterProfiler v3.18.1, colorspace v2.1-0, DOSE v3.28.0, dplyr v1.0.7, enrichplot v1.22.0, ggplot2 v3.3.5, GOplot v1.0.2, jsonlite v1.7.2, limma v3.58.1, orgDbs (like org.Hs.eg.db v3.16.0), stringr v1.4.0, tidyverse v1.3.1 |
|  | DiffGO-GSEA | Same to the analysis function of DiffGO |
|  | DiffKEGG | KEGG pathway database, clusterProfiler v3.18.1, colorspace v2.1-0, DOSE v3.28.0, dplyr v1.0.7, enrichplot v1.22.0, ggplot2 v3.3.5, GOplot v1.0.2, jsonlite v1.7.2, limma v3.58.1, orgDbs (like org.Hs.eg.db v3.16.0), stringr v1.4.0, tidyverse v1.3.1 |
|  | DiffKEGG-GSEA | Same to the analysis function of DiffKEGG |
|  | DiffTreePlot | Same to the analysis function of DiffGO |
|  | Pathview | Same to the analysis function of DiffKEGG |
| CorrExpToolkit | CorrExp | dplyr v1.0.7, jsonlite v1.7.2 |
|  | PairedCorr | dplyr v1.0.7, ggplot2 v3.3.5, ggpubr v0.4.0 |
|  | UnpairedCorr | dplyr v1.0.7, ggplot2 v3.3.5, ggpubr v0.4.0 |
|  | Corrplot | corrplot v0.89, dplyr v1.0.7, ggplot2 v3.3.5 |
|  | CircosPlot | circlize v0.4.15, corrplot v0.89 |
|  | NetworkPlot | igraph v1.5.1, reshape2 v1.4.4 |
|  | CorrGO | Same to the analysis function of DiffGO |
|  | CorrKEGG | Same to the analysis function of DiffKEGG |
| WGCNAToolkit | WGCNA | clusterProfiler v3.18.1, dplyr v1.0.7, ggplot2 v3.3.5, jsonlite v1.7.2, scatterplot3d v 0.3-41, stringr v1.4.0, WGCNA v 1.70-3 |
|  | WGCNA GO | Same to the analysis function of DiffGO |
|  | WGCNA KEGG | Same to the analysis function of DiffKEGG |
| FeatureSelectToolkit | COX Regression | dplyr v1.0.7, survival v3.2-11, survminer v0.4.9, tidyr v1.3.0 |
|  | Survival | dplyr v1.0.7, survival v3.2-11, survminer v0.4.9, tidyr v1.3.0, TSHRC v0.1.6 |
|  | LASSO | dplyr v1.0.7, glmnet v4.1-3, foreign v0.8-81, jsonlite v1.7.2, |
|  | ROCCurve | ggpubr v0.4.0, plotROC v2.2.1, randomcoloR v1.1.0.1, RColorBrewer v1.1-3, ROCR v1.0-11, tidyverse v1.3.1 |
